# Filtering input fluctuations in intensity and in time underlies transcriptional pulses without feedback

**DOI:** 10.1101/2020.03.24.006445

**Authors:** Alberto Stefano Sassi, Mayra Garcia-Alcala, Philippe Cluzel, Yuhai Tu

## Abstract

Stochastic pulsatile dynamics have been observed in an increasing number of biological circuits with typical mechanism involving feedback control. Surprisingly, recent single-cell experiments showed that *E. coli* flagellar class-2&3 promoters are activated in stochastic pulses without the means of feedback, however, the underlying design principles of pulse generation have remained unclear. Here, by developing a system-level stochastic model constrained by a large set of *E. coli* flagellar synthesis data from different strains and mutants, we identify the underlying design principles for generating stochastic transcriptional pulses without feedback. Our model shows that YdiV, an inhibitor of the class-1 master regulator (FlhDC), creates an ultrasensitve switch that serves as a digital filter to eliminate small amplitude FlhDC fluctuations. Additionally, we demonstrate that fast temporal fluctuations of FlhDC are smoothed out and integrated over time before affecting class-2 downstream genes. Together, our results reveal the existence of a filter-and-integrate design that is necessary for generating stochastic pulses without feedback. This strategy suggests that *E. coli* may avoid premature activation of the expensive flagellar gene expression by filtering input fluctuations in intensity and in time.

## I. INTRODUCTION

The gene regulatory network for controlling the bacterial flagellar synthesis and assembly has long served as a canonical example of transcriptional cascade [1–8], in which a particular gene product (protein) can serve as the transcription factor (TF) for other genes, which encode proteins that serve as TF for other downstream genes, and so on. The bacterial flagellar promoters are organized in three classes, each of which underlies specific stages of the complex flagellar assembly and chemo-sensory signaling processes [9–11]. The class-1 promoter controls the transcription of *flhDC*, the operon encoding for the proteins FlhD and FlhC, that assemble into the heterohexamer FlhD_4_C_2_ (refer to as FlhDC hereafter), which is the master regulator of flagellar synthesis. The master regulator serves as the TF for class-2 promoters and initiates the transcription of seven operons necessary for the assembly of the basal body and the hook of the flagellum [12, 13]. A specific class-2 protein, FliA, activates class-3 promoters to express class-3 genes coding proteins for the flagellar filament and the chemotaxis signaling pathway [14]. The bacterial flagellar synthesis represents a remarkable process in which over 50 different genes are expressed in a coherent fashion over several generations to build a large functional protein complex. Over the past decades, experimental studies have yielded many valuable insights about the transcriptional network that underlies flagellar synthesis in *E. coli* and *Salmonella* [9, 10, 15–18]. In particular, population measurements suggested that the transcriptional cascade follows a deterministic temporal order where genes in subsequent classes are activated sequentially and once turned on promoters remain active continuously during exponential growth [10].

Recently, Kim et al. [19] measured gene expression dynamics of single *E. coli* cells over many generations by using a microfluidic device called the “mother machine” [20, 21]. Surprisingly, the new single-cell data showed that flagellar promoters are stochastically activated in intermittent pulses, i.e., bursts of strong activity amid a quiet background. Due to their stochastic nature, transcriptional pulses can only be detected by single-cell measurements and they were missed in previous cell population study [10].

Indeed, rapid developments in quantitative single-cell measurements have revealed that quasi-periodic or stochastic pulsating dynamics is a common dynamic activity pattern in biological circuits in a wide range of organisms, e.g., transcription factor Msn2 in yeast [22], transcription factor p53 [23] and NF-*κ*B [24] in mammalian cells, bacterial transcription factor *σ*_*b*_ in the bacterium Bacillus [25], and stochastic cell fate switching in microbes [26]. Typically, the underlying mechanism involves a combination of positive and negative feedback loops [27], a recent theoretical study showed that stochastic pulses may also be caused by a negative feedback loop alone [28]. However, in Kim et al [19] pulses were present even when the promoter of FlhDC was deleted and replaced by a synthetic/constitutive promoter that does not respond to flagellar endogenous regulator and thus rules out the involvement of any feedback control. Additionally, in a different system that controls cell fate, stochastic pulses were also successfully generated by transferring in *E. coli* a reconstituted gene circuit without feedback [29].

Prompted by these recent observations, we ask what are the underlying design principles that govern the observed stochastic transcriptional pulses in flagellar synthesis without feedback? To answer this question, we use the large single-cell expression time series dataset for different strains and mutants from Kim et al [19] to quantitatively and consistently constrain a system-level stochastic model. Using this strategy, we aim to identify the minimum design principles necessary to generate stochastic transcriptional pulses without feedback during flagellar biosynthesis and in general.

## II. RESULTS

We first present a system-level (coarse-grained) stochastic model for the regulation of class-2 genes by FlhDC. We then use the model to analyze and explain the steady-state distributions of class-2 activity and its dynamics observed in single-cell experiments [19]. Next, we propose possible molecular mechanisms underlying the main findings from our analysis and describe a filter-and-integrate mechanism for controlling noisy FlhDC signal in *E. coli*. Finally, we use our model to make testable predictions for responses to realistic time-varying signals.

### A. A system-level stochastic model for class-1 and class-2 gene expression dynamics

In the experiments b Kim et al [19], they used strains whose expression of class-1 genes was controlled by promoters of different strengths (see Sec. A in Supplemental Information (SI) for details of the experimental data). For each strain, long time series (∼ 70 generations) of class-1 and class-2 promoter activity was simultaneously measured within the same cells using fluorescent proteins as proxy. They found that distributions of class-2 promoter activity depend on the class-1 promoter strength as well as the presence or absence of the inhibitor YdiV [30–32] (also known as RflP [33]).

To be consistent with the experiments, we define the class-2 gene expression activity, *A*_2_, as the time derivative of the fluorescence time series of the class-2 reporter gene, and use the class-1 reporter florescence signal as a proxy to represent the FlhDC level *C*_1_. We propose a system-level stochastic model to describe dynamics of these two coarse-grained observables, *A*_2_ and *C*_1_, without considering molecular details. In particular, the *A*_2_ dynamics in a single cell is described by a stochastic differential equation (SDE) that has three basic terms: decay (degradation), activation (production), and noise:

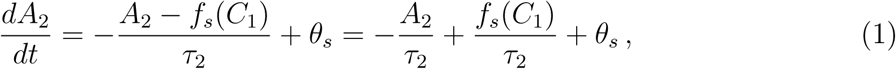

where *s* = (+, −) represents cells with and without YdiV respectively, and 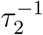 is the decay rate with *τ*_2_ the characteristic timescale associated with class-2 expression. For convenience, we write activation rate of the class-2 promoter as *f*_*s*_(*C*_1_)/*τ*_2_ so that the average 〈*A*_2_〉 = 〈*f*_*s*_(*C*_1_)〉. We call *f*_*s*_(*C*_1_) the response function as it characterizes how class-2 activity responds to class-1 protein level *C*_1_. The white noise *θ*_*s*_ has zero mean *〈θ*_*s*_(*t*)〉 = 0 and correlation *〈θ*_*s*_(*t*)*θ*_*s*_(*t′*)〉 = 2∆_*s*_(*〈C*_1_〉)*δ*(*t* − *t′*) with noise strength given by ∆_*s*_(*〈C*_1_*〉*). Here, we assume that the noise strength ∆_*s*_(*〈C*_1_*〉*) depend only on the average *C*_1_ concentration *〈C*_1_*〉*, which is valid due to the timescale difference for *A*_2_ and *C*_1_ as will become clear later in this work.

The activity of class-2 genes *A*_2_ depends on the class-1 protein level *C*_1_, whose dynamics can be modeled by a similar SDE:

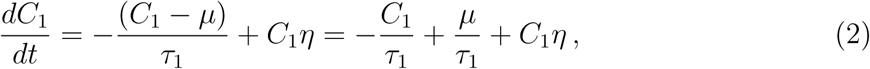

where *τ*_1_ is the decay time for class-1 protein. We define *µ*/*τ*_1_ as the *C*_1_ production rate so that the steady-state average *〈C*_1_*〉* = *µ* with *µ* depending on the promoter strength of a specific strain. The white noise *η* has zero mean *〈η*(*t*)*〉* = 0 and *〈η*(*t*)*η*(*t′*〉) = 2∆_1_*δ*(*t* − *t′*) with the noise strength ∆_1_ a constant for all promoter strengths. The noise *η* is multiplied by *C*_1_ in Eq. 2. This choice of noise comes directly from class-1 reporter fluorescence intensity measurement data, which can be better fitted by having a multiplicative noise than an additive noise in Eq. 2 (see Sec. B in SI for details).

Overall, our “minimal” system-level (coarse-grained) model consists of two SDE’s (Eqs. 1–2) with a few physiologically meaningful parameters and functions: two timescales *τ*_1_ and *τ*_2_, a constant multiplicative noise strength ∆_1_, a strain-dependent noise strength ∆_*s*_(*µ*), and two response functions *f*_*s*_(*C*_1_). In the following, by fitting all the single-cell data for different strains [19] with our model quantitatively, we determine the parameters and response functions, which help us identify the underlying mechanism for transcriptional pulses.

### B. YdiV creates an ultrasensitive switch

We first use our model to study all the steady state class-2 activity distributions from experiments (see Methods for details on simulations of our model). In Fig. 1A, experimentally observed distributions of the normalized class-2 activity *A*_2_*/〈A*_2_*〉* for cells with and without YdiV for three class-1 promoter strengths: low (P1, red), medium (P4, green), and high (P7, blue) are shown (see Sec. A in SI for details of the experimental dataset). The most drastic difference between cells with and without YdiV is for strain P4 that has a medium promoter strength similar to that of the wild-type cells. The *A*_2_*/〈A*_2_*〉* distribution for the P4 strain in the presence of YdiV (green line in the left panel of Fig. 1A) is strongly asymmetric with a peak that is significantly shifted with respect to the mean, whereas the distributions for lower and higher promoter strength are more symmetric. This asymmetry is largely absent in strains without YdiV (the right panel of Fig. 1A). Our model can explain all the steady state *A*_2_ distributions from different strains together as shown by the direct comparison between experimentally measured distributions (Fig. 1A) and those obtained from our model (Fig. 1B).

**FIG. 1:**
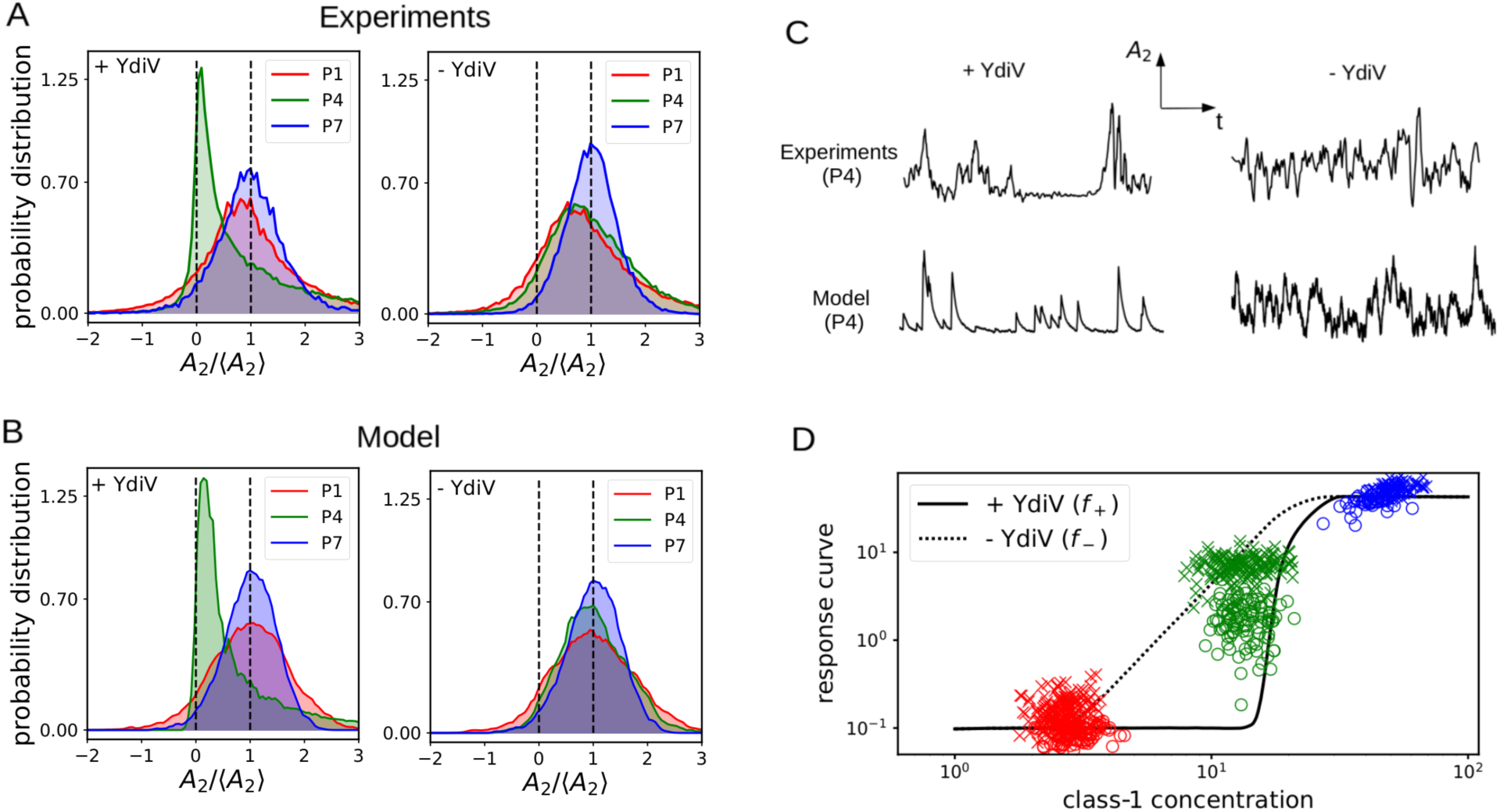
Comparison between experimental data and model results. (A) Distributions of the normalized class-2 activity 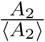 in the presence (left) and absence (right) of YdiV for small (P1, red), medium (P4, green) and large (P7, blue) class-1 promoter strengths. Data taken from [19]. We use the black dashed lines to indicate the positions of its mean value and zero for *A*_2_. (B) The corresponding distributions obtained from our model. (C) Comparison of typical time series of *C*_1_(*t*) and *A*_2_(*t*) from experiments and model for the P4 strain with and without YdiV. (D) Response functions obtained by fitting our model to experimental data [19] with (*f*_+_, solid line) and without YdiV (*f*_−_, dashed line). The circles (+YdiV) and crosses (-YdiV) indicate the mean value from different mother cells in experiments. Note that *f*_*s*_ does not pass through the center of the data points due to its high nonlinearity (*〈f*_*s*_(*C*_1_)*〉* ≠ *f*_*s*_(*〈C*_1_*〉*). The color code for each promoter is the same as in A&B.

The most significant difference in the dynamics of class-2 activity *A*_2_(*t*) between cells with and without YdiV also occurs in strain P4 with a medium class-1 promoter strength similar to that of wild-type cells. As shown in Fig. 1C, from the experiments, the *A*_2_ dynamics contains well-defined pulses in the presence of YdiV, while pulses are not well separated and the presence of higher frequency noise appears to be unfiltered in the absence of YdiV. These different activity dynamics are reproduced in our model.

By fitting our model to the single-cell data (see Sec. C in SI for details of the fitting and values of all the fitted model parameters), we obtain the response functions, *f*_+_(*C*_1_) and *f*_−_(*C*_1_), which cannot be directly determined from experimental data because *〈A*_2_*〉* = *〈f*_*s*_(*C*_1_)*〉* ≠ *f*_*s*_(*〈C*_1_*〉*) due to their nonlinearity. As shown in Fig. 1D (solid line), the response function *f*_+_(*C*_1_) in the presence of YdiV behaves like a ultrasensitive switch, i.e., it has a small constant value up to a threshold where it rises sharply. Due to this ultrasensitive (steep) response function, when the mean FlhDC concentration is close to the threshold (e.g., for strain P4), the distribution of *A*_2_*/〈A*_2_*〉* has the largest asymmetry and there is a significant shift in the peak of the distribution (left panels in Fig. 1A&B). In the absence of YdiV, the response function *f*_−_(*C*_1_) is found to be less steep (dotted line in Fig. 1D) and the probability distributions are more symmetric (right panels in Fig. 1A&B). The agreements between our model and the experiments not only valid our model, more importantly, they reveal that the role of YdiV is to create a ultrasensitive (steep) response of the class-2 activity to the class-1 protein (FlhDC) concentration.

### C. Separation of timescales and memory effect

Next, we use our model to analyze the stochastic gene expression dynamics in the flagellum system. As shown in Eq. 1, there are two noise sources for the class-2 activity *A*_2_: the intrinsic noise *θ*_*s*_ in the gene expression process and the extrinsic noise due to fluctuations of the FlhDC concentration *C*_1_. From our model, the mean class-2 promoter activity averaged over the intrinsic noise can be written in the following integral form:

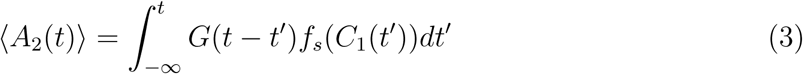

where 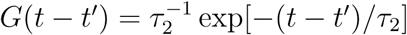 is the integral kernel (Green’s function) that decays with the timescale *τ*_2_. Eq. 3 means that *A*_2_(*t*) depends on the values of *C*_1_(*t′*) in a time window *t* − *τ*_2_ ≤ *t′* ≤ *t*, i.e., there is a memory effect. Given that *C*_1_(*t*) varies with a timescale *τ*_1_, the strength of this memory effect depends on the ratio of the two timescales *τ*_2_/*τ*_1_. For *τ*_2_/*τ*_1_ « 1, the kernel *G*(*t* − *t′*) ≈ *δ*(*t* − *t′*) and *〈A*_2_(*t*)*〉* ≈ *f*_*s*_(*C*_1_(*t*)), i.e., the class-2 activity depends on the instantaneous level of class-1 protein *C*_1_(*t*) with negligible memory effect. However, if *τ*_2_/*τ*_1_ » 1, the memory effect is dominant as 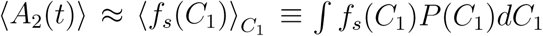 becomes a constant independent of the instantaneous FlhDC level *C*_1_(*t*) (*P* (*C*_1_) is the steady state *C*_1_ distribution function).

To demonstrate the memory effect, we divide the FlhDC concentration in equally spaced bins (in log-scale) and then compute the class-2 activity and FlhDC concentration averaged within each bin to form the so called “bin-averaged response curve” (BARC) for each strain with a different class-1 promoter strength (see Methods for details on BARC). As shown in Fig. 2A, when *τ*_2_ > *τ*_1_, the BARC’s for different strains can be shifted from each other consistent with experiments. This means that the class-2 activity is not determined just by the instantaneous FlhDC concentration, it depends also on its average over previous times due to the memory effect. Indeed, as shown in Fig. 2B, in the absence of memory effect when *τ*_2_ « *τ*_1_, all BARC’s collapse.

**FIG. 2:**
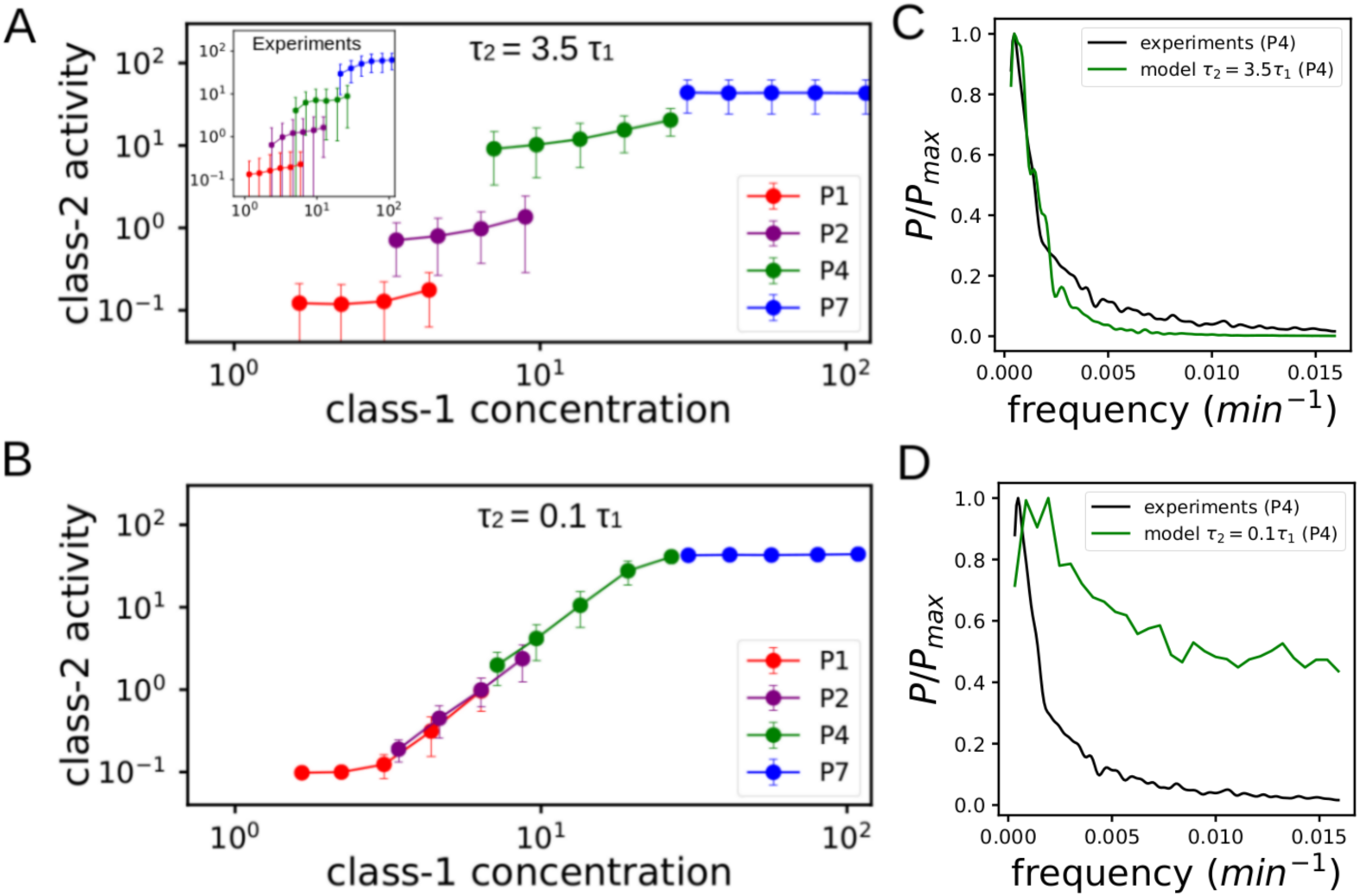
The bin-averaged response curves and the ratio of two timescales *τ*_2_/*τ*_1_. (A) Model results for *τ*_2_ = 3.5*τ*_1_. The inset shows the same behavior from experimental data. (B) Model results for *τ*_2_ = 0.1*τ*_1_. we used -YdiV strains here, see SI for similar results for +YdiV strains. The normalized power spectra (green lines) from our model for *τ*_2_ = 3.5*τ*_1_ and *τ*_2_ = 0.1*τ*_1_ for the P4 strain are shown in (C)&(D), respectively. The normalized power spectrum (black line) from experiments (P4) is also shown for comparison. We used *τ*_1_ = 45 *min.* within the range of *C*_1_ correlation time estimated from experiments.

When *τ*_2_ > *τ*_1_, *A*_2_ depends on an integral of the transformed input signal *f*_+_(*C*_1_) over a time window *τ*_2_ as shown in Eq. 3. This integration effectively averages the input signal in time and eliminates high frequency fluctuations as demonstrated in Fig. 2C&D where power spectra for the two cases with and without integration are shown. The power spectrum from experimental data is consistent with our model with *τ*_2_ = 3.5*τ*_1_ where the power of the spectrum of *A*_2_(*t*) is concentrated in the low frequency regime with a similar half-maximum-power frequency *f*_1/2_ ≈ 0.002*/min* as shown in Fig. 2C. For *τ*_2_ « *τ*_1_ (Fig. 2D), the power spectrum is much broader with significant high frequency fluctuations and *f*_1/2_ ≈ 0.008*/min* that is much higher than that from experiments. Quantitatively, by comparing BARC’s and autocorrelation functions from the model and those from experiments, we estimate the timescale ratio to be in the range: 2.1 ≤ *τ*_2_/*τ*_1_ ≤ 3.6 (see Sec. D in SI for details).

### D. A YdiV-mediated sequestration mechanism for filtering

So far, we have used our model to explain the steady state distributions and dynamics of the noisy single-cell data. In this section, we propose possible molecular mechanisms underlying our findings with a focus on the role of YdiV in regulating FlhDC signal.

We first briefly discuss possible molecular origins of separation of timescales. The shorter timescale *τ*_1_ is most likely related to the degradation time of the FlhC, and FlhD monomers and the longer timescale *τ*_2_ is related to the decay time of the FlhDC-dependent class-2 promoter activation (see Fig. 3A for an illustration). Since FlhD&FlhC can only serve as functional transcription factor when they form heterohexamer FlhDC, *τ*_2_ is affected by the binding affinity of FlhDC to class-2 promoters as well as stability of the FlhDC complex. The heterohexamer is more stable than the monomers [34], which is consistent with *τ*_2_ > *τ*_1_ as observed in experiments and our model. It would be interesting to test the effects of changing *τ*_2_ experimentally (see the Discussion section for more details).

**FIG. 3:**
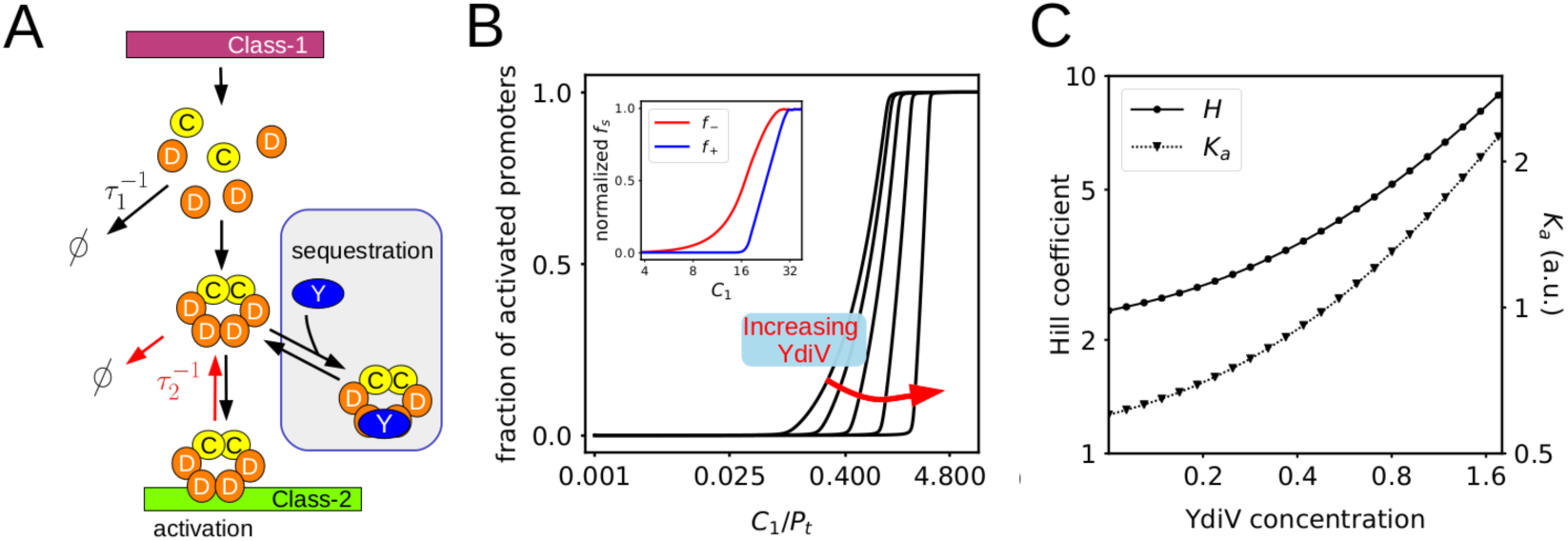
The YdiV-mediated sequestration mechanism for switch like behavior. (A) Schematic representation of the mechanism of transcriptional regulation of class-2 promoters by FlhDC. The class-1 promoter (purple) initiates the expression of the monomers FlhC and FlhD (yellow and orange respectively) that forms the heterohexamer FlhD_4_C_2_ (FlhDC). The timescale *τ*_1_ may be related to the degradation rate of the monomers (FlhD and FlhC), while *τ*_2_ may be related to the degradation rate of FlhDC or its unbinding rate to the promoters, both labeled by red arrows. The activation of class-2 genes, which depends on binding of the FlhDC complex to the class2 promoter (green), is inhibited when FlhDC is sequestered by binding to YdiV (blue) instead (gray box). (B) Response curves obtained from the YdiV sequestration model for different values of YdiV concentrations (*P*_*t*_ is the total class-2 promoter concentration), see Methods for details. (inset) the response functions, *f*_+_(*C*_1_) (blue) and *f*_−_(*C*_1_) (red), in the presence and absence of YdiV respectively from fitting experimental data (the same as in Fig. 1D). (C) Hill coefficient (*H*) and the half maximum concentration (*K*_*a*_) as a function of the YdiV concentration.

We now study the role of YdiV in creating an ultrasensitive switch-like response of class-2 gene expression as shown in Fig. 1D. YdiV is known to have two main effects on FlhDC. First, by occupying the binding sites of FlhDC, YdiV prevents the binding of FlhDC to free class-2 promoters [31]. Second, YdiV mediates the interaction between FlhDC and the degradation complex ClpXP [30, 31]. Here, we only consider the effect of competitive binding on the response function, similar effect may be achieved by enhancing the degradation of FlhDC, which is not considered in this study. Intuitively, due to competitive binding, YdiV can introduce a threshold for FlhDC by sequestrating FlhDC to prevent it from binding to and activating the class-2 promoters.

To test this YdiV-mediated sequestration mechanism, we developed a model based on competitive binding of FlhDC by YdiV and the class-2 promoter as illustrated in Fig. 3A (see Methods for details of the sequestration model). In Fig. 3B, the fraction of promoters bound to FlhDC, which serves as a measure of the class-2 gene expression activity, is shown as a function of the scaled FlhDC concentration *C*_1_/*P*_*t*_, with *P*_*t*_ the total promoter concentration, for different values of total YdiV concentration. We can see that higher YdiV concentrations lead to delayed but steeper response curves, which agrees qualitatively with the response curves *f*_+_ and *f*_−_ shown in the inset of Fig. 3B for comparison. A response curve can be quantitatively described by its Hill coefficient *H* and its half maximum concentration *K*_*a*_. From the experimental data, the Hill coefficient changes roughly from *H* ∼ 2 in the ∆*ydiV* mutant to *H* ∼ 4 in the wild type, and *K*_*a*_ almost doubles. As shown in Fig. 3C, the YdiV-mediated sequestration model reproduces the general trend with both *H* and *K*_*a*_ increasing with YdiV concentration in a range consistent with experiments.

The steep response function *f*_+_(*C*_1_) suggests that YdiV serves (approximately) as a digital filter. When the input signal, i.e., FlhDC concentration (*C*_1_) is lower than a threshold *C*∗ ≡ *K*_*a*_(*Y*_*t*_) set by the YdiV concentration *Y*_*t*_, the normalized output *f*_+_(*C*_1_)*/max*(*f*_+_) is close to 0. When *C*_1_ > *C*∗, the normalized output is close to 1. The (normalized) filtered signal switches between 0 and 1 randomly resembling a random telegraph signal.

### E. A filter-and-integrate mechanism for stochastic pulses without feedback

Put together, our results show that *E. coli* uses a combined filter-and-integrate strategy (mechanism) to control its class-2 gene expression. As shown in Fig. 4, in the presence of YdiV (left side of Fig. 4), the input signal, i.e., the FlhDC concentration *C*_1_(*t*) (blue line), which fluctuates relatively fast at a short timescale *τ*_1_, first goes through a highly nonlinear ultrasensitive response function *f*_+_(*C*_1_), which serves approximately as a digital filter that eliminates the FlhDC fluctuations lower than certain threshold *C*∗ set by the YdiV concentration. The filtered signal *f*_+_(*C*_1_(*t*)) (green line) has a nearly digital stochastic pulse-like pattern akin to the random telegraph signal.

**FIG. 4:**
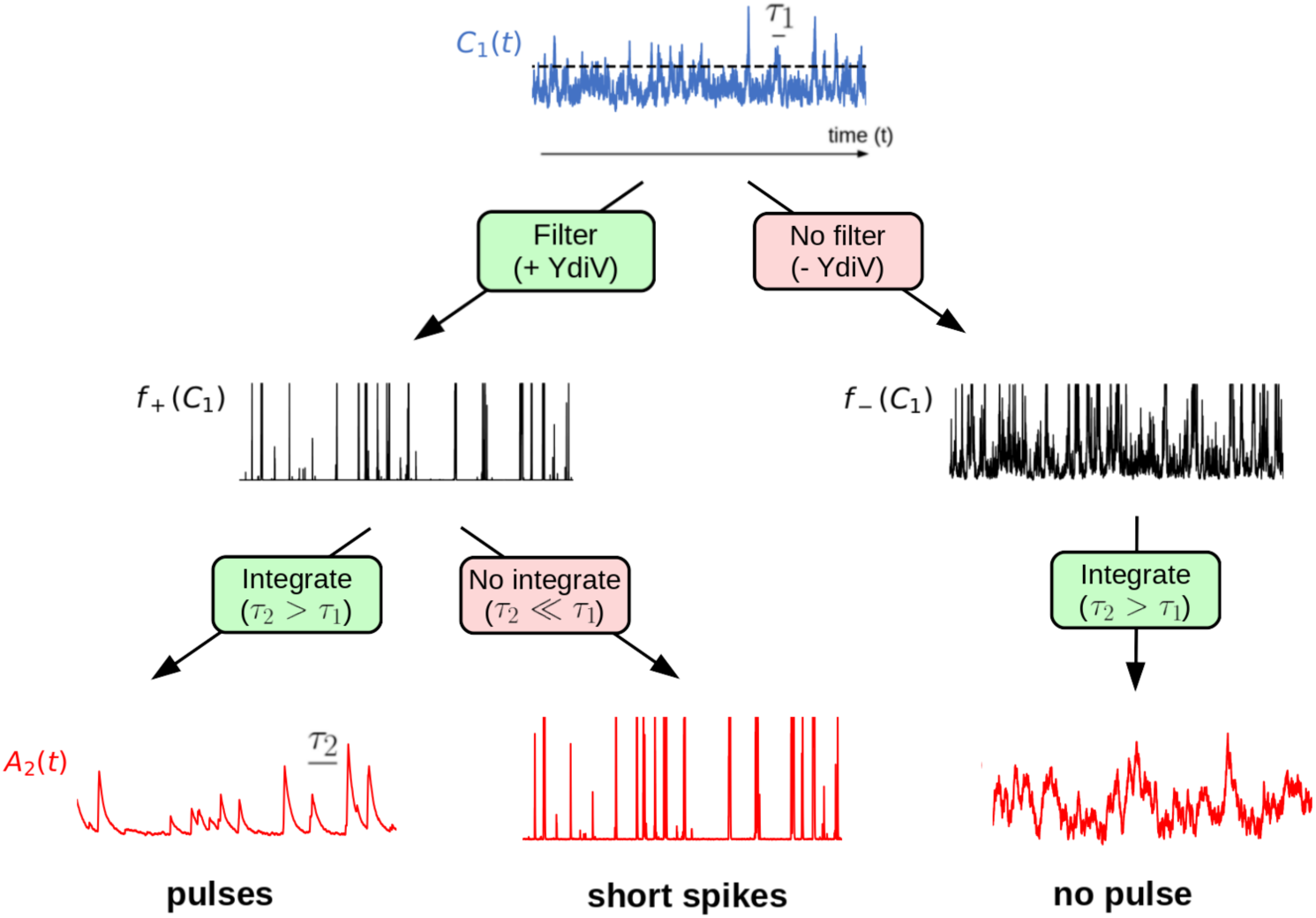
The filter-and-integrate mechanism for pulse generation. In the presence of YdiV, the FlhDC concentration *C*_1_(*t*) (blue line), which fluctuates with a fast timescale *τ*_1_, is filtered by the inhibitory effect of YdiV. The filtered signal *f*_+_(*C*_1_(*t*)) (black line), which has a pulse-like pattern, is then integrated over a longer timescale *τ*_2_ to determine the class-2 activity *A*_2_(*t*) (red line, bottom left). In the absence of YdiV, there is no filtering effect, and the resulting *A*_2_ dynamics is noisier with no pulse (bottom right). Without integration (*τ*_2_ « *τ*_1_), *A*_2_ would have many short spikes (bottom middle). Results presented here are obtained from simulations of our model using parameters fitting the P4 strain.

The second part of the two-pronged process is to integrate the filtered signal over a timescale *τ*_2_(> *τ*_1_), which gives rise to the intermittent “strong” pulses in *A*_2_ (red line). Without the integration process *τ*_2_ « *τ*_1_, the *A*_2_ activity would have many short spikes, too short to complete the flagellar synthesis process. The integration process acts as a filter in frequency space as demonstrated in Fig. 2C&D to suppress those FlhDC fluctuations that are strong in intensity but short in time duration. As a result, class-2 gene expression is turned on only when there is persistently high levels of FlhDC signal. Due to the same integration (memory) effect, once turned on, the class-2 expression remains active for a time duration ∼ *τ*_2_ even without strong instantaneous FlhDC signal. This long duration of ontime is necessary for the cell to finish the flagellar synthesis process that lasts for several generations.

Both filtering and integration in time are critical to generate the stochastic pulses observed in experiments [19] where *A*_2_ alternates between having near zero activity (quiet) for a long period of time and being highly active during a strong pulse that lasts for a long timescale set by *τ*_2_ (bottom left in Fig. 4). In the absence of YdiV, the noisy *C*_1_(*t*) signal is not filtered and as a result there is no pulse in *A*_2_(*t*), which fluctuates continuously (bottom right in Fig. 4). In the absence of integration, *A*_2_ follows the same dynamics as the filtered signal *f*_+_, which has many short spikes (bottom middle in Fig. 4).

### F. Model predictions: gene expression responses to time-varying signals

Now that our model is fully developed with all its parameters determined by existing single-cell experiments [19], it provides a powerful tool to predict class-2 gene expression responses to more realistic time-dependent signals without any tuning parameter. As an example, we use our model to study the responses of class-2 activity to an increase of the class-1 promoter strength from a low value *µ*_0_(= *µ*(*P*_1_) ≈ 2.6) to a series of higher values *µ*_1_ at time *t* = 0. We find that a cell will turn on the class-2 gene expression after a delay time *τ*_*r*_ as shown in Fig. 5A. The delay time *τ*_*r*_ varies from cell to cell and follows a distribution *P* (*τ*_*r*_|*µ*_1_) that depends on *µ*_1_. In Fig. 4B, the delay time distributions for different values of *µ*_1_ are shown. Both the average delay time and its variance decrease with *µ*_1_ or equivalently the average FlhDC level as shown in Fig. 4C. Responses to other time-varying stimuli such as ramps and oscillatory signals can be studied by our model similarly.

**FIG. 5:**
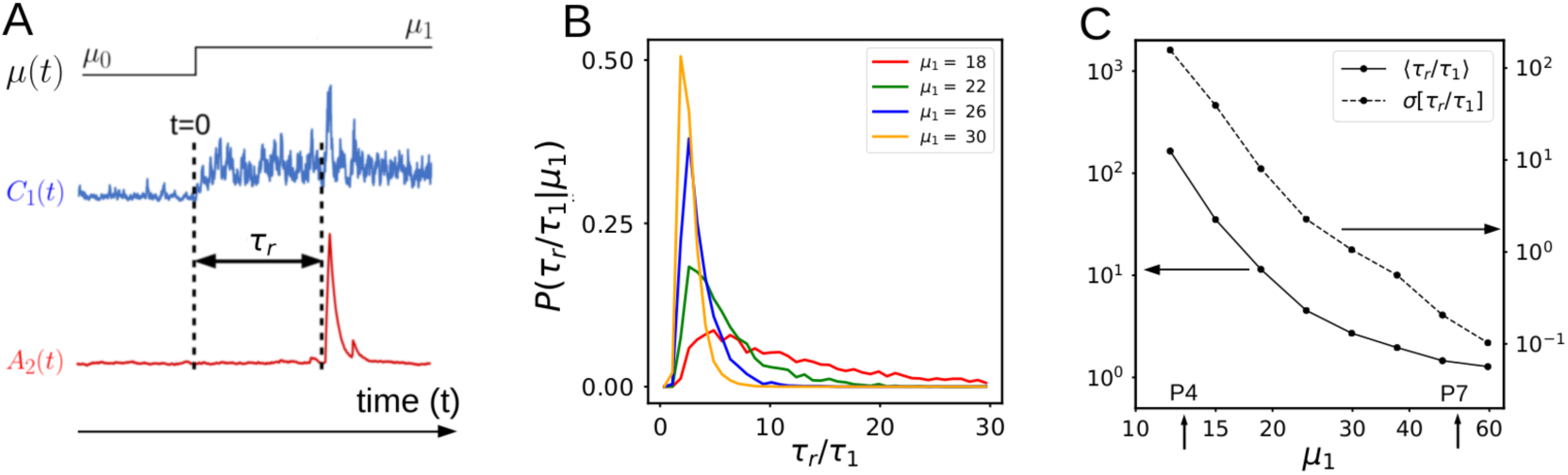
The predicted responses to step stimuli. (A) Predicted dynamics of *C*_1_ and *A*_2_ in response to a time-dependent (step function) stimulus in which the class-1 promoter strength *µ* is changed at *t* = 0 from a low value *µ*_0_ = *µ*(*P*1) = 2.6 to a higher value *µ*_1_. The delay time *τ*_*r*_ is the time duration from the stimulus (*t* = 0) to the onset of class-2 expression, which is defined as the time when *A*_2_ cross a threshold 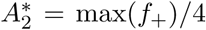 (qualitative results do not depend on the choice of 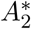). (B) Distributions of the delay time *τ*_*r*_ (normalized by *τ*_1_) for different values of *µ*_1_. (C) Both the average and variance of *τ*_*r*_ decreases with *µ*_1_ or the average FlhDC concentration. The values of *µ* for strains *P*4 and *P*7 used in our study are marked for reference.

Our results suggest that even though FlhDC concentration does not control the class2 gene expression deterministically, the filter-and-integrate mechanism allows the cell to control the timing statistics of the class-2 expression based on the average FlhDC signal intensity. As shown in Fig. 4C, a higher FlhDC concentration induces (on average) a shorter and more accurate (smaller variance) onset time to turn on the class-2 genes. Similarly, an elevated average FlhDC level leads to more frequent occurrence of the transcriptional pulses. These model predictions can be tested in future single cell experiments by using inducible class-1 promoters.

## III. SUMMARY AND DISCUSSIONS

In this paper, dynamics of the class-1 and class-2 promoters in *E. coli* flagellar synthesis are studies by using a minimal stochastic model that captures the essential characteristics of the underlying system. Applying our stochastic model to a large single-cell dataset quantitatively shows that the transcriptional pulses are generated by a filter-and-integrate mechanism without the need of any feedback control. Both the YdiV-enabled filtering and the memory-enabled time integration are crucial to average out FlhDC fluctuations in intensity and in time so that an individual cell can make a “calculated” (informed) decision on whether to turn on the expensive class-2 gene expression. The same general mechanism may be at work in other pulse-generating systems without feedback, e.g., in the reconstituted SinI-SinR circuit [29], the antagonist SinI may play the role of the filter (analogous to YdiV) by binding to and inhibiting the transcriptional regulator SinR (analogous to FlhDC).

To better understand the feedback-less filter-and-integrate strategy in *E. coli*, we compare it with a closely related bacterium *Salmonella*, which does have an additional positive feedback loop in controlling its class-2 promoter activity. As illustrated in Fig. 6, FliZ, a class-2 protein, can inhibit YdiV in *Salmonella* but not in *E. coli*. This positive feedback loop in *Salmonella* leads to bistability in class-2 gene response [35]. As a result, the hysteretic response function in *Salmonella f*_*Sal*_(*C*_1_) has two critical points 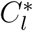 and 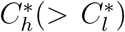 as illustrated in Fig. 6B. Due to the bistability, once the FlhDC level goes over the upper critical level 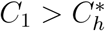 even for a short time, the response signal *f*_*Sal*_(*C*_1_) will stay on (green) for a much longer time as long as FlhDC level is higher than the lower critical level 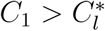. This amplification of the “on”-time duration allows *Salmonella* to activate class-2 gene expression upon detecting a large-but-short signal. In contrast, for *E.coli* as shown in Fig. 6A, the response function, though steep, does not have bistability because there is no feedback. As a result, an elevated FlhDC level over the threshold (*C*∗) for a short time is not enough to trigger significant class-2 activity given the large integration time *τ*_2_.

**FIG. 6:**
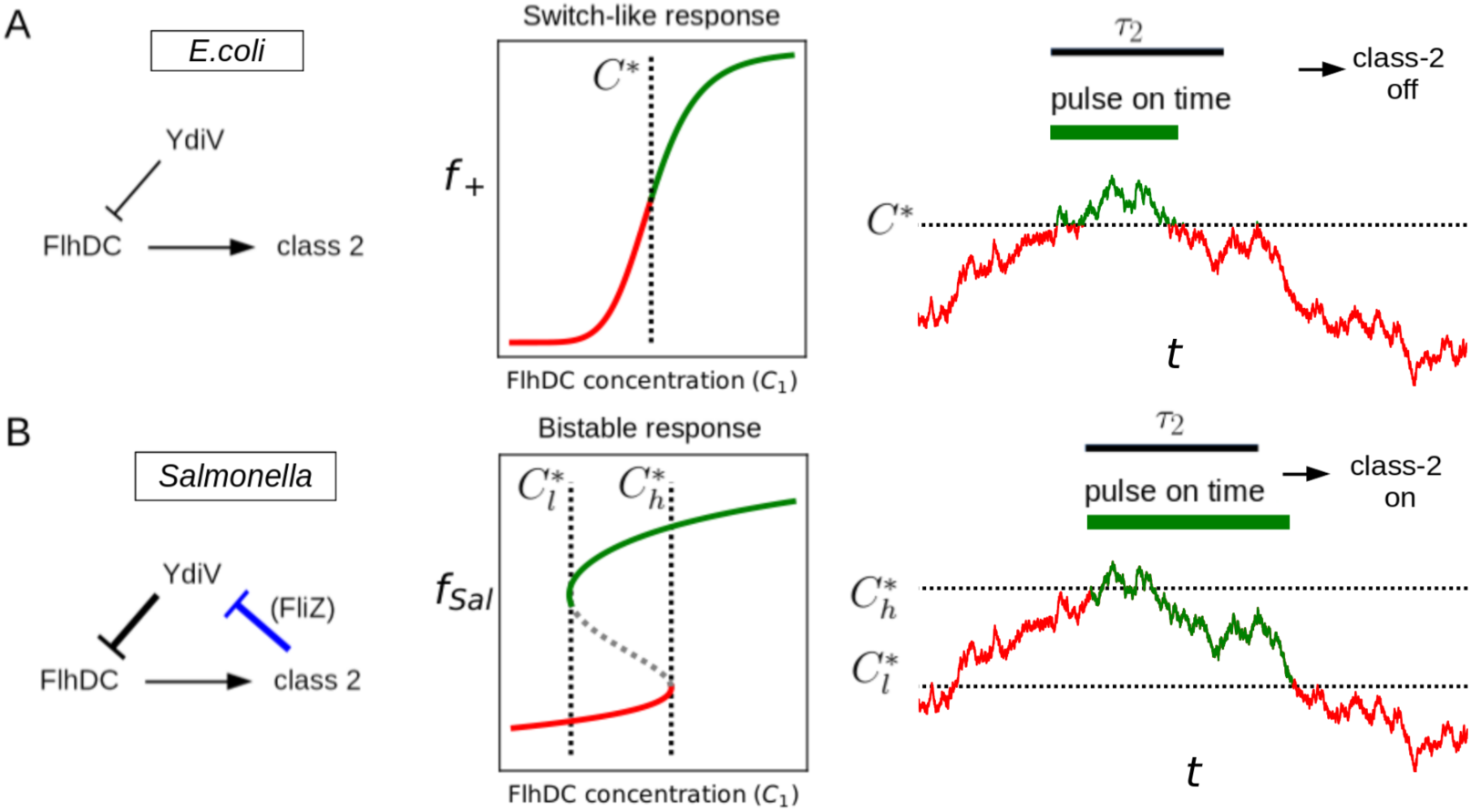
The different gene regulation strategies for *E. coli* and *Salmonella*. (A) The *E. coli* network has the YdiV-mediated inhibition of the class-1 master regulator (FlhDC) but not the feedback mechanism from class-2 gene. As a result, *E. coli* has a steep but monotonic response function *f*_+_ with a threshold *C*∗, i.e., class-2 gene expression is activated (green) when *C*_1_ > *C*∗ or inhibited (red) when *C*_1_ < *C*∗. For a FlhDC signal that passes the threshold *C*∗ briefly, the class-2 activity is only turned on for a short time, which is not enough to start the flagella synthesis given the long integration time *τ*_2_ for class-2 gene activation. (B) In *Salmonella*, there is a feedback (blue line) from a class-2 protein (FliZ) to suppress the inhibitor YdiV. This feedback mechanism leads to a bistable (hysteretic) response of class-2 genes in the range 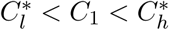. As a result, a brief period of elevated FlhDC concentration over the (upper) threshold 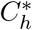> can cause a prolonged class-2 gene activation as long as 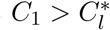, which can lead to flagella synthesis.

Their different strategies may reflect the different purposes of motility for these two bacteria. *Salmonella* is a pathogen and activates the flagellar gene expression cascade in nutrient rich conditions. A short pulse of strong stimulus from the host may trigger the flagellar synthesis process. This aggressive strategy may be optimized for host colonization and infection. In *E. coli*, however, flagellar synthesis is activated in nutrient poor conditions for the purpose of foraging and searching for nutrients. Therefore, *E. coli* may adopt a more prudent (filter-and-integrate) strategy that only triggers the expensive flagellar synthesis process when the cell experiences persistent poor nutrient conditions.

Our current analysis opens up a few new directions for future investigation. On the experiment side, effects of the filter-and-integration mechanism may be tested by perturbing the two key factors, e.g., by tuning the strength of the interaction between FlhDC and class-2 promoters [31, 36] or by reducing the effective concentration of YdiV by adding *Salmonella* FlhC known to have weaker affinity for *E. coli* promoters but the same affinity for YdiV as *E. coli* FlhC [37]. It would be highly informative to measure single-cell gene expression dynamics of *Salmonella* in the mother machine as done for *E. coli* [19] to understand its gene regulation strategy quantitatively. On the modeling side, YdiV level is assumed to be constant in our model as we focus on the long timescale (low frequency) dynamics of the system. However, in the high frequency regime, the *A*_2_ power spectrum from experiments follows *P* (*f*) ∼ *f*^−1^ (*f* is the frequency), which decays slower than the power spectrum from our current model with a fixed YdiV level (see Fig. S7 in SI). Similar 1/*f* noise spectrum was observed in flagellar motor switching dynamics [38] and was subsequently explained by considering fluctuations of the response regulator CheY-P in a single cell [39]. It would be interesting to include dynamics of YdiV in our model to explore whether it also leads to the observed 1*/f* spectrum in the high frequency regime.

Finally, our work here demonstrates that system-level stochastic modeling augmented by quantitative single-cell measurements for different strains and mutants provides a powerful tool to decipher principles of gene regulation from inherently noisy single-cell data. This tool becomes necessary when dealing with complex gene regulatory circuits such as the full bacterial flagellar gene expression cascade with multiple feedback controls. The general system-level stochastic modeling approach with coarse-grained variables that can be directly compared with experiments should be applicable to study stochastic expression dynamics in other gene regulatory systems with or without feedback [22–25, 29, 40].

## IV. METHODS

### Simulation of the stochastic differential equations

For the numerical simulation of the stochastic equations, we write Eq. 2 in discrete form:

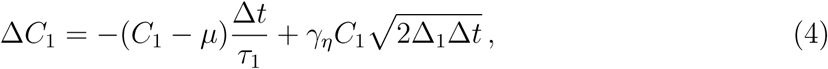

where *γ*_*η*_ is a stochastic variable following a Gaussian distribution 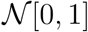. We let *τ*_1_ = 1 to set the timescale, and the time step ∆*t* = 0.004*τ*_1_. The discrete form of Eq. 1 is:

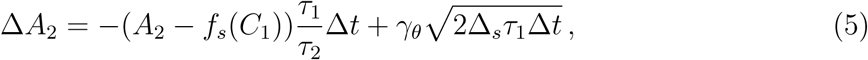

where *γ*_*θ*_ is a Gaussian random number as *γ*_*η*_.

### Bin-averaged response curves and flatness

To obtain the bin-averaged response curves, we divide the full interval 10^−1^ ≤ *C*_1_ ≤ 10^2^ into 20 logarithmically spaced bins and calculate the average of class-1 concentration and class-2 activity for all the data points in each bin. This bin-averaged class-2 activity versus the bin-averaged class-1 concentration is called the bin-averaged response curve (BARC). The BARC in Fig. 2 are only shown at those bins that contain more than 5000 data points for better statistics.

### Steady-state solution of the sequestration model

Let *P*_*t*_ be the concentration of promoters, *C*_1_ is the total concentration of FlhDC, *Y*_*t*_ the YdiV concentration. We denote *C*_1*p*_ and *C*_1*y*_ as the concentrations of FlhDC that are bound to the promoters and YdiV respectively. In steady state, the balance of binding and unbinding reactions leads to: 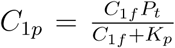, and 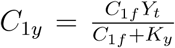, where *K*_*p*_ and *K*_*y*_ are the dissociation constants of FlhDC to the promoters and YdiV respectively. *C*_1*f*_ (= *C*_1_ − *C*_1*p*_ − *C*_1*y*_) is the free FlhDC concentration and it satisfies the following equation (conservation of FlhDC):

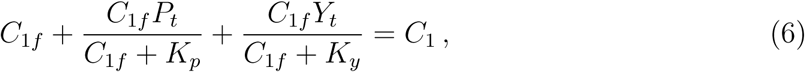

which can be solved to determine *C*_1*f*_ as a function of *P*_*t*_, *C*_1_, *Y*_*t*_, and *K*_*y*_, *K*_*p*_. The fraction of FlhDC bound promoters is *C*_1*p*_/*P*_*t*_ = *C*_1*f*_/(*K*_*p*_ + *C*_1*f*_), which is used as a measure of the class-2 activity. The dependence of *C*_1*p*_/*P*_*t*_ on *C*_1_/*P*_*t*_ is plotted in Fig. 3B. Parameters *K*_*y*_/*P*_*t*_ = 10^−2^ and *K*_*p*_/*P*_*t*_ = 10^−1^ are used in Fig. 3B&C.

## V. ACKNOWLEDGMENTS

We thank Drs. M. Kim and M. Erhardt for discussions. The work by ASS and YT are supported by a NIH grant (R35GM131734 to YT). The work by MG and PC are supported by a NIH grant (R01GM134275 to PC) and a NSF grant (1615487 to PC).

## Supplementary Information

### A: The single-cell experimental dataset

All the experimental data analyzed in this paper are from the published single cell experiments by Kim et al [19]. Briefly, to vary the class-1 expression strength, different constitutive artificial promoters are used to replace the native class-1 promoter (flhDp) and either T7 or mut4 sequence are used as the downstream ribosome binding site (RBS-1). The ratio of the effective number of class-1 proteins and fluorescent proteins produced is different for the mut4 and T7 strains. To account for this difference, we multiply the *C*_1_ values by a factor (∼ 0.22) for the strains with mut4 as RBS-1 (see SI in [19] for details).

The experimental dataset consists of ∼ 100 independent single-cell time series for each class-1 promoter strain, each lasting for ∼ 70 generations. The time interval between two consecutive time frames is 5 minutes. Given that the average division time is ∼ 40 minutes, each time series has ∼ 70 × 40/5 = 560 time points on average, and there are on average ∼ 100 × 560 = 56, 000 data points for each strain. In this paper, we consider four of the seven strains studied in [19]: P1, P2, P4 and P7 (we use the same strain names as in [19]). These four strains are chosen to cover the full range (∼ 20-fold) of class-1 promoter strength change in [19].

### B: Distribution of class-1 concentration and multiplicative noise

In order to model the class-1 protein concentration we used a multiplicative noise in Eq. 2 in the main text rather than an additive noise. This choice is motivated by the form of the probability distribution. We can consider for example the class-1 distribution when the promoter strength is P4 (green data points in Fig. S1). In case of additive noise, the concentration follows an Ornstein-Uhlenbeck process (OU) whose steady state distribution is Gaussian. In Fig. S1, we plot a Gaussian distribution (black curve) with the same mean and variance as the experimental data as well as the distribution (red curve) obtained with a multiplicative noise. Fig. S1 clearly shows that the fluctuations in *C*_1_ are better described by a stochastic model (Eq. 2 in the main text) with a multiplicative noise.

### C: Model parameters and response functions determined by fitting model to the single-cell experimental dataset

In this paper, we develop a simple stochastic model with a minimal set of biologically meaningful parameters and response functions, which can be determined by fitting our model to the large experimental data set described in Sec. A.

For the *C*_1_ dynamics described in Eq. 2 in the main text, there are three parameters: the timescale *τ*_1_, the multiplicative noise strength ∆_1_, and the promoter strength *µ*, which is different for different strains. In our study, *τ*_1_ can be set to unity to set the timescale of the problem, all other timescales are scaled by *τ*_1_. By fitting our model to the experimental data, we found that the noise strength 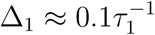, and the promoter strength for different strains are: *µ*(*P*1) = 2.6, *µ*(*P*2) = 6.06, *µ*(*P*4) = 13.5, *µ*(*P*7) = 51.7, in the unit of florescence intensity.

The dynamics of class-2 activity *A*_2_ is determined by a timescale *τ*_2_, the response function *f*_*s*_(*C*_1_), and a noise strength ∆_*s*_ that is different for different strains. The parameters and response functions are obtained by fitting our model to the experimental data (class-2 activity dynamics). The range of *τ*_2_/*τ*_1_ is determined as shown in Fig. 2 in the main text. The noise strength ∆_+_ and ∆_−_ for different strains in the presence and absence of YdiV are also determined and their values are shown in Fig. S2.

For the response function *f*_*s*_ (with *s* = +, −), we parameterize it with a piece wise power law function that approximate a sigmoid function with a low and a high plateau. In particular, we set a few anchor points (*x*_*i*_, *y*_*i*_) (*i* = 1, 2,.., *N*) for *f*_*s*_ such that *f*_*s*_(*x*_*i*_) = *y*_*i*_. Between the anchor points *x*_*i*_ ≤ *x* ≤ *x*_*i*+1_, 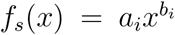 follows a power law, and we have *f*_*s*_(*x* ≤ *x*_1_) = *y*_1_ and *f*_*s*_(*x* ≥ *x*_*N*_) = *y*_*N*_ to enforce the two plateaus. We determine the coordinates of the anchor points by fitting our model (with the piece wise power law response function) to the data. For the response function *f*_+_ in the presence of YdiV, we used *N* = 4 anchor points with their fitted coordinates: *f*_+_(*C*_1_ ≤ 17) = 0.1, *f*_+_(19) = 0.2, *f*_+_(22) = 10, *f*_+_(*C*_1_ ≥ 24) = 43. For the response function *f*_+_ in the absence of YdiV, we used only *N* = 2 anchor points with their fitted coordinates: *f*_−_(*C*_1_ ≤ 3) = 0.1 and *f*_−_(*C*_1_ ≥ 22) = 43.

In Fig. S3, we show the average *C*_1_ versus average *A*_2_ for a single cell over many generations from experiments and model. The response functions (smoothed around the anchor points) are also shown (the same as in Fig. 1D in the main text).

### D: Estimating the timescale ratio

The separation of timescales seems to be independent of the presence or absence of YdiV. In Fig. S4, we show the BARC’s from experimental data for strains with YdiV and those from our model, which are similar to the results for strains without YdiV (Fig. 2 in the main text). Quantitatively, BARCs can be used to estimate the timescale ratio *τ*_2_/*τ*_1_. To quantify the shape of BARC, we define a flatness parameter as the absolute value of the inverse of the average slope of linear fits to BARC’s of different strains such as those shown in Fig. 2 in the main text and Fig. S4 here. A larger flatness means that the class-2 activity has a stronger dependence on the average FlhDC level than its instantaneous value and thus the memory effect is stronger. We computed the flatness for different values of the relative timescale ratio *τ*_2_/*τ*_1_ in our model. As shown in Fig. S5, a larger *τ*_2_/*τ*_1_ ratio leads to a larger flatness, i.e., a stronger memory effect. Based on a quantitative comparison with the flatness values obtained from experimental data with and without YdiV, we estimate the ratio *τ*_2_/*τ*_1_ to be in the interval, 2.1 ≤ *τ*_2_/*τ*_1_ ≤ 3.6, shown as the gray region in Fig. S5. In addition, in Fig. S6, we compare the autocorrelation function for wild type cells shown in [19] with the autocorrelation functions obtained from the simulations of our model (Eq. 1 and 1). The difference in the timescales between class-1 and class-2 is in line with the interval estimated by the analysis of the flatness shown in Fig. S5.

---

**FIG. S1:**
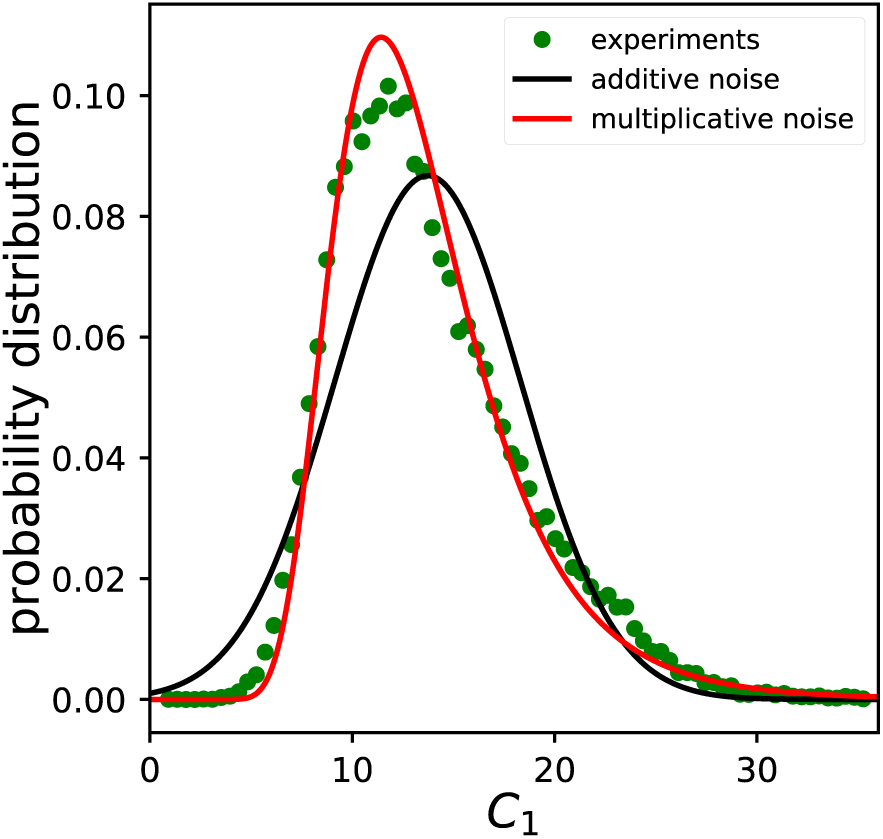
Comparison of the model with additive and multiplicative noise to fit the experiments. In the case of multiplicative noise the values are distributed following an inverse Gamma function (red). This curve fits the experimental distribution (green) more accurately than the Gaussian distribution (black), corresponding to a model with additive noise.

**FIG. S2:**
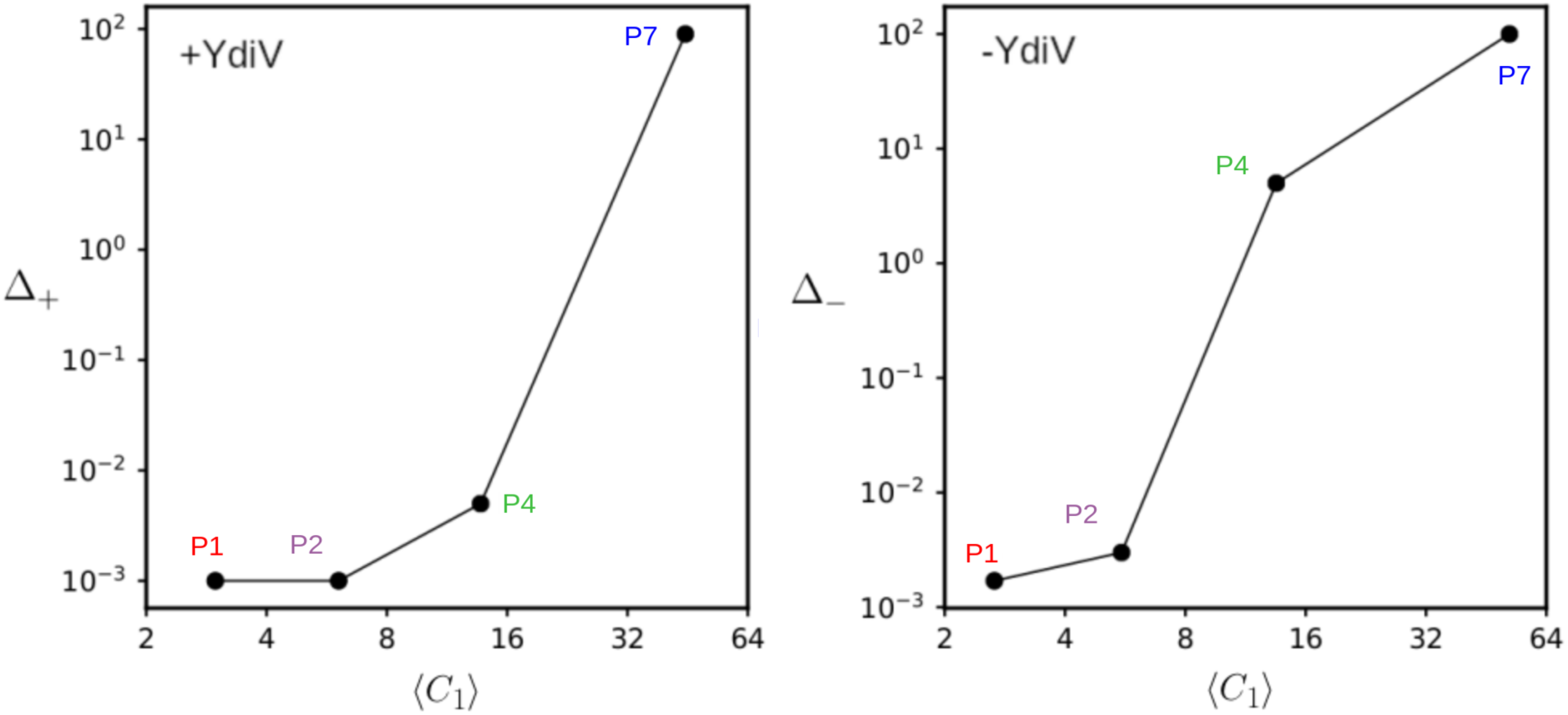
Noise strength ∆_+,−_ (in the presence and absence of YdiV) as a function of the average class-1 protein level 〈*C*_1_〉 obtained by fitting the stochastic model (Eq. (1–2) in the main text) to experimental data.

**FIG. S3:**
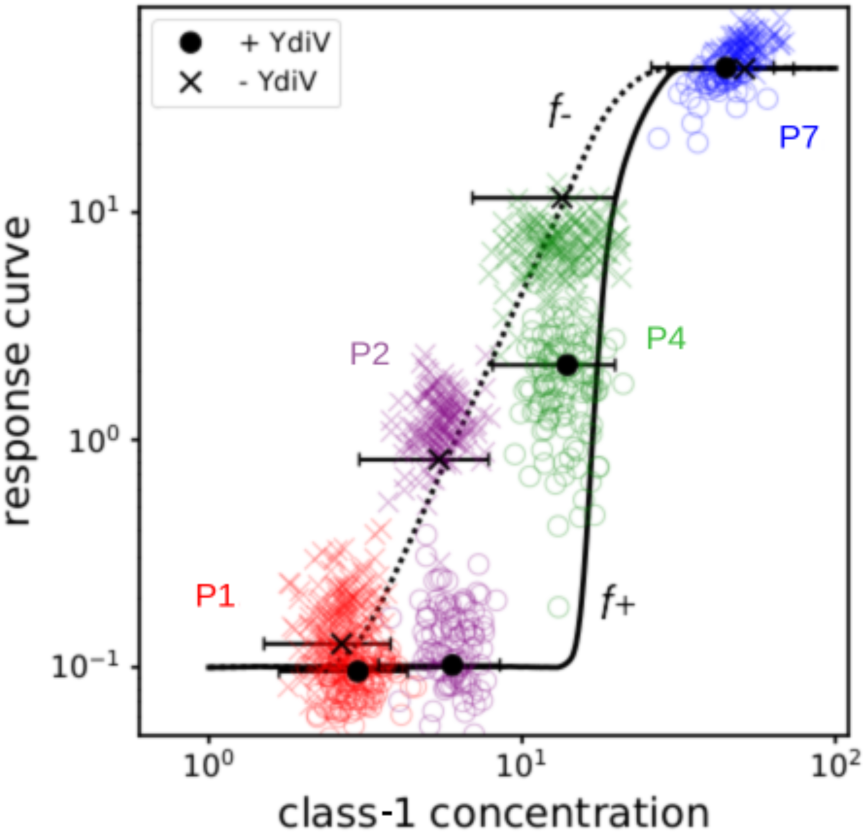
Response functions *f*_+,−_(*C*_1_) used in the model, in the presence (solid) and absence (dashed) of YdiV. The black circles and crosses indicate the mean value obtained with the stochastic model. The horizontal error bars (for *C*_1_) are given by 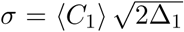, where ∆_1_ is the strength of the noise *η* for class-1 protein level.

**FIG. S4:**
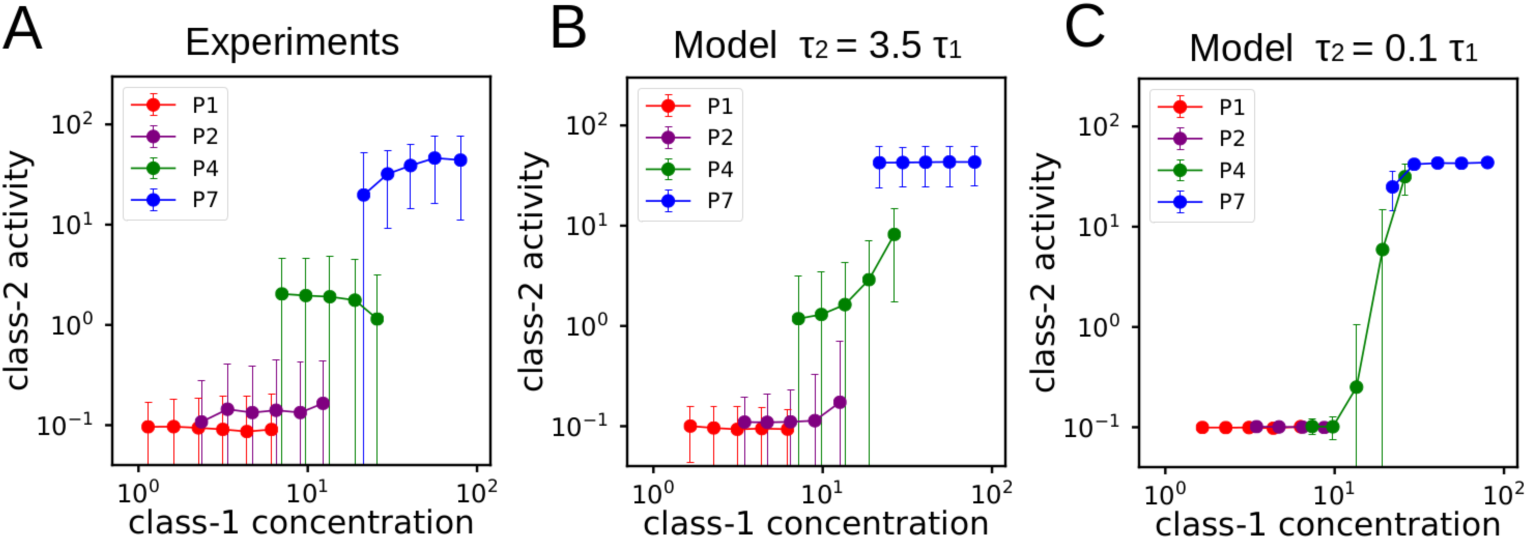
The bin-averaged response curves and the ratio of two timescales *τ*_2_*τ*_1_. (A) The bin-averaged response curves with different class-1 promoter strength from experimental data [19] for strains with YdiV. Different colors are used for different class-1 promoters. The corresponding model results for (B) *τ*_2_ = 3.5*τ*_1_ and (C) *τ*_2_ = 0.1*τ*_1_.

**FIG. S5:**
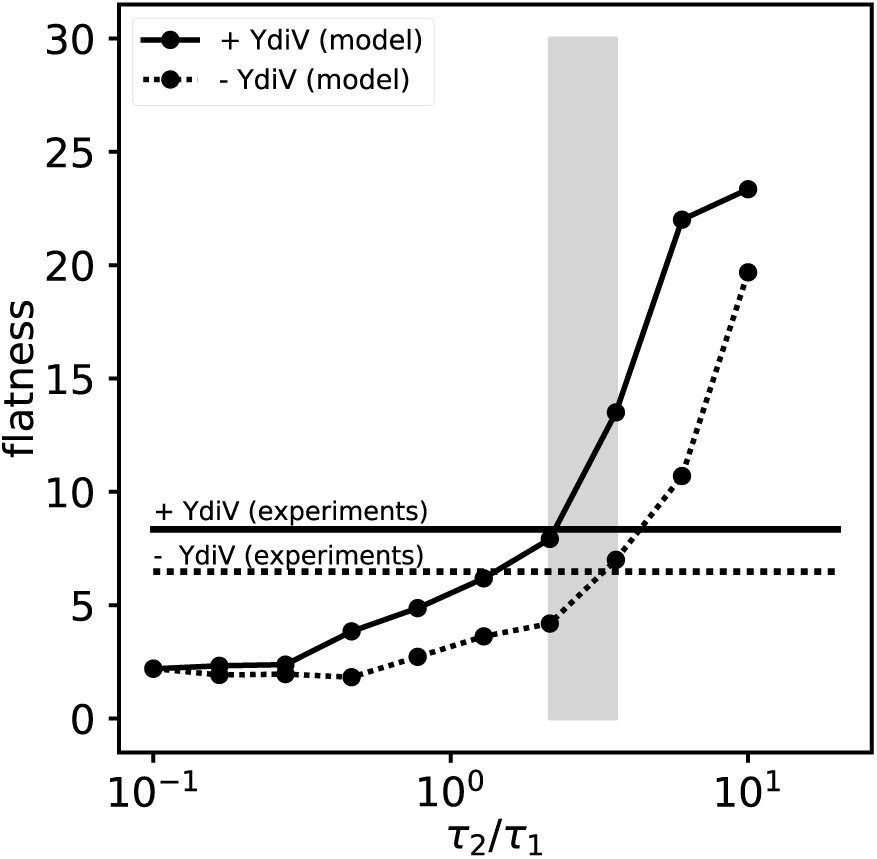
The estimated range of the timescale ratio *τ*_2_/*τ*_1_. Flatness of the bin-averaged response curves as a function of *τ*_2_/*τ*_1_ in the absence (dashed line) and presence (solid line) of YdiV. The horizontal lines indicate the experimental results. Comparing the model results with the experiments, we estimate the ratio *τ*_2_/*τ*_1_ to be within the gray region: 2.1 ≤ *τ*_2_/*τ*_1_ ≤ 3.6.

**FIG. S6:**
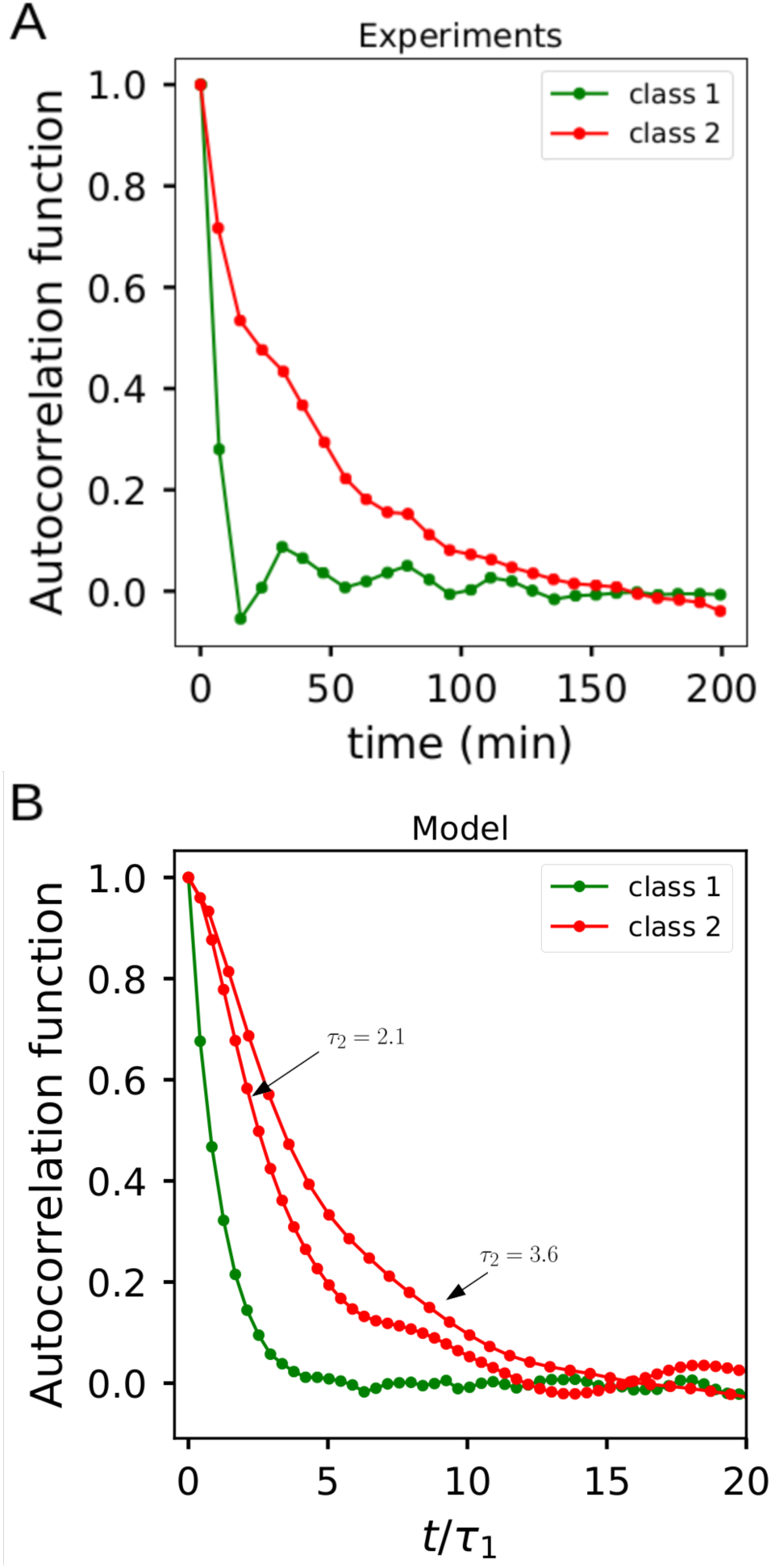
(A) Autocorrelation functions for the fluorescence of class-1 and class-2 reporters in the case of wild type promoters (taken from Fig. 2D in [19]). (B) Autocorrelation functions for *C*_1_ and *A*_2_ from the stochastic simulations of our model (Eq.1 and Eq.2) for P4 strain.

**FIG. S7:**
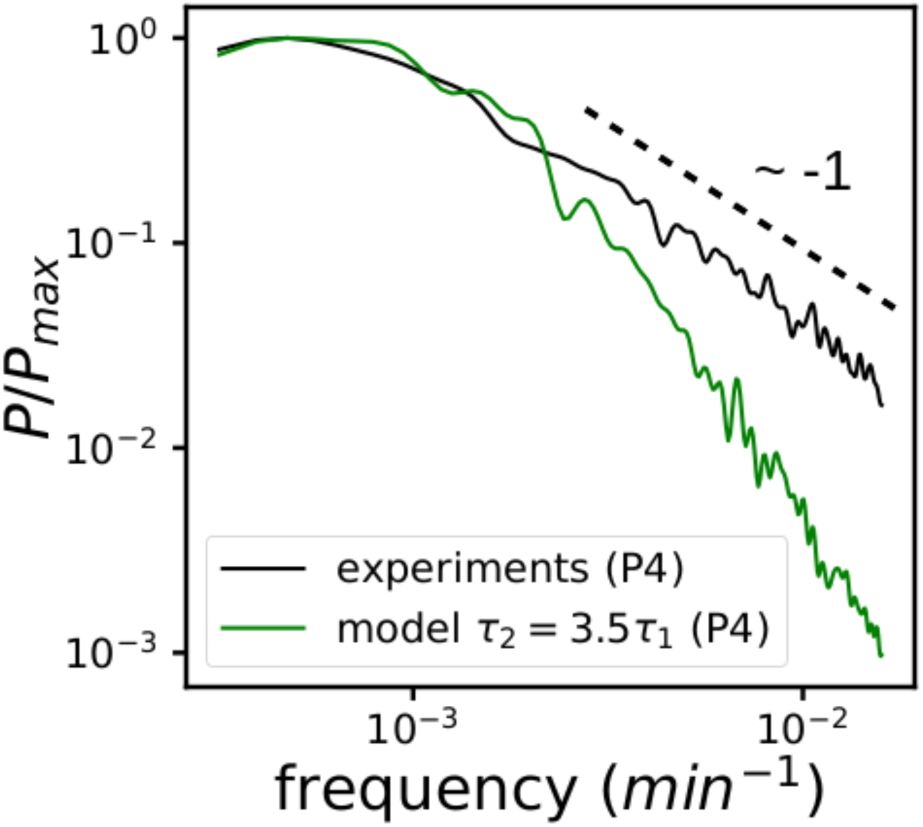
The power spectrum in log-log scale. The power spectrum from experimental data agrees with our model in the low frequency regime. In the high frequency regime, it decays as *f*^−1^ slower than the model result. This 1/*f* noise may be caused by fluctuations of YdiV level, which can introduce a spectrum of timescales to the system.

## Notes

### Competing Interest Statement

The authors have declared no competing interest.

### Summary of Updates

Several changes have been made to the text in order to stress the importance of the absence of feedback in the mechanism described. Figures with the power spectrum of the class-2 activity from model and experiments have been added.

